# Neuronal overexpression of mouse potassium channel subunit Kcnn1 in A53T α-synuclein mice more than doubles median survival time, associated with suppression of phospho-S129 α-synuclein formation

**DOI:** 10.64898/2026.03.09.709927

**Authors:** Maria Nagy, Wayne A. Fenton, Arthur L. Horwich

## Abstract

Synucleinopathies, including idiopathic Parkinson’s Disease, are driven by misfolding and aggregation of the 140 residue α-synuclein protein that plays a role in presynaptic vesicle regulation. We describe effects of a modifier, neuronal overexpression of the mouse calcium-activated potassium channel subunit Kcnn1, on a mouse model in which transgenic Thy1.2-driven A53T α-synuclein directs fully penetrant lethal motor disease. Kcnn1 overexpression increased median survival of these mice from 8.5 months to 18 months, associated with an altered clinical presentation from a rapidly progressive dystonic-like behavior of the limbs to a later-onset (12-16 mo) and slowly progressive lower limb clasping when lifted by the tail. At the tissue level, accretion of disease-associated phospho-serine 129 α-synuclein was prevented by overexpression of Thy1.2-driven Kcnn1, which was observed in many brain regions, including the ones where phospho-serine 129 α-synuclein was copiously accreted in A53T mice at endstage. The action of blocking production of phospho-serine 129 α-synuclein was also observed in adult presymptomatic A53T mice injected with an AAV9 scCMV-Kcnn1 virus into the right superior colliculus. At endstage ∼2 months later, the right superior colliculus exhibited overexpression of Kcnn1 and showed essentially no phospho-serine 129 α-synuclein, whereas the uninjected left superior colliculus exhibited copious phospho-serine 129 α-synuclein. The neuroprotective action of Kcnn1 overexpression remains to be fully resolved, but the channel protein subunit, targeted to the ER membrane, has been shown to induce an ER stress response. This response, which may activate autophagy, along with potential channel formation, may diminish the rate of formation or lifetime of neurotoxic forms of A53T α-synuclein.

---

Idiopathic Parkinson’s Disease is one of several neurodegenerative disorders called synucleinopathies, in which inclusions called Lewy bodies and Lewy neurites are formed, composed principally of filaments of aggregated α-synuclein, an abundant 140 residue protein of neuronal tissue that normally appears to play a role in presynaptic vesicle regulation.^1–6^ In the diseased state, α-synuclein assembles into oligomeric^7^ and fibrillar^8^ forms that exert toxic behavior, leading to neuronal dysfunction and loss. Clinically, by mid-stage of Parkinson’s Disease, this prominently affects the nigrostriatal dopaminergic system, leading to a classic movement disorder of tremor/rigidity/bradykinesia/postural instability^1^. Further spreading of pathologic forms of α-synuclein within the brain can produce broader effects including dementia^9^.

Rare pedigrees with early-onset autosomal dominant-inherited Parkinson’s Disease have been found to harbor mutations in the α-synuclein gene, e.g. A53T^10^ or A30P^11^, further implicating α-synuclein in Parkinson’s Disease. These mutant synucleins were subsequently programmed as neuronal-overexpressed transgenes in mice, producing α-synuclein aggregation in brain and spinal cord, associated with various motor symptoms (see refs 12 and 13 for review). The brain regions affected correlated significantly with the neuronal promoter employed and its regions of greatest expression. The nigrostriatal dopaminergic system was generally spared but several strains were affected to varying degree^14–16^. Additionally, formation of disease-associated phospho-serine 129-α-synuclein was reported in brains of model strains^17,18^. While the mouse models have various imperfections, they allow for study of aspects of α-synuclein pathogenesis and testing of therapeutic strategies.

Here, we have further examined a genetic modifier effect on an A53T mutant α-synuclein transgenic mouse strain. Transgenic-programmed neuronal overexpression of Kcnn1, the subunit of a calcium-activated mouse potassium channel, served to more than double the median survival time of the strain. This was associated with absence of disease-associated phospho-serine 129 α-synuclein. Similarly, overexpression of Kcnn1 from intracerebral-delivered AAV9-CMV-Kcnn1, injected into a superior colliculus of adult presymptomatic A53T mice, suppressed formation of phospho-serine 129 α-synuclein observed at endstage.

## RESULTS

### Extended survival of double transgenic A53T α-synuclein/Kcnn1-6copy mouse strain

As reported previously^19^, a Thy1.2 neuronal promoter-driven human A53T α-synuclein cDNA transgene, introduced into a B6SJL background at ∼20 copies, produced fully penetrant lethal motor disease, with a median survival of 8½ months^19^. Mice commenced forward flexion/splaying of lower extremities at 6-7 months, developed a wobbling gait with sudden lurches sideways, and then at late time assumed dystonic posturing involving splaying of all four extremities followed by inability to right (see ref.19 Supp. Fig. 11), requiring sacrifice. At endstage, copious phosphoserine 129 α-synuclein was observed in a number of regions e.g. motor cortex, deep cerebellar nuclei, zona incerta, spinal cord. We crossed in a transgenic modifier (from a B6SJL background) that we had previously observed to extend survival of SOD1-linked ALS mice^19^, the Thy1.2-driven potassium channel subunit Kcnn1 cDNA, at both 3 copies and 6 copies (separate transgene insertions). We had observed that at 3 copies the median survival of A53T mice was extended to 12½ months (∼45%)^19^. We have now followed a cohort of the A53T/Kcnn1-6copy mice and observed that the median survival was extended to 18 months (vs 8½ mo for A53T), comprising greater than 100% extension of median survival (Fig.1, green trace vs black, p = 1.6 x 10^-10^).

**Figure 1.**
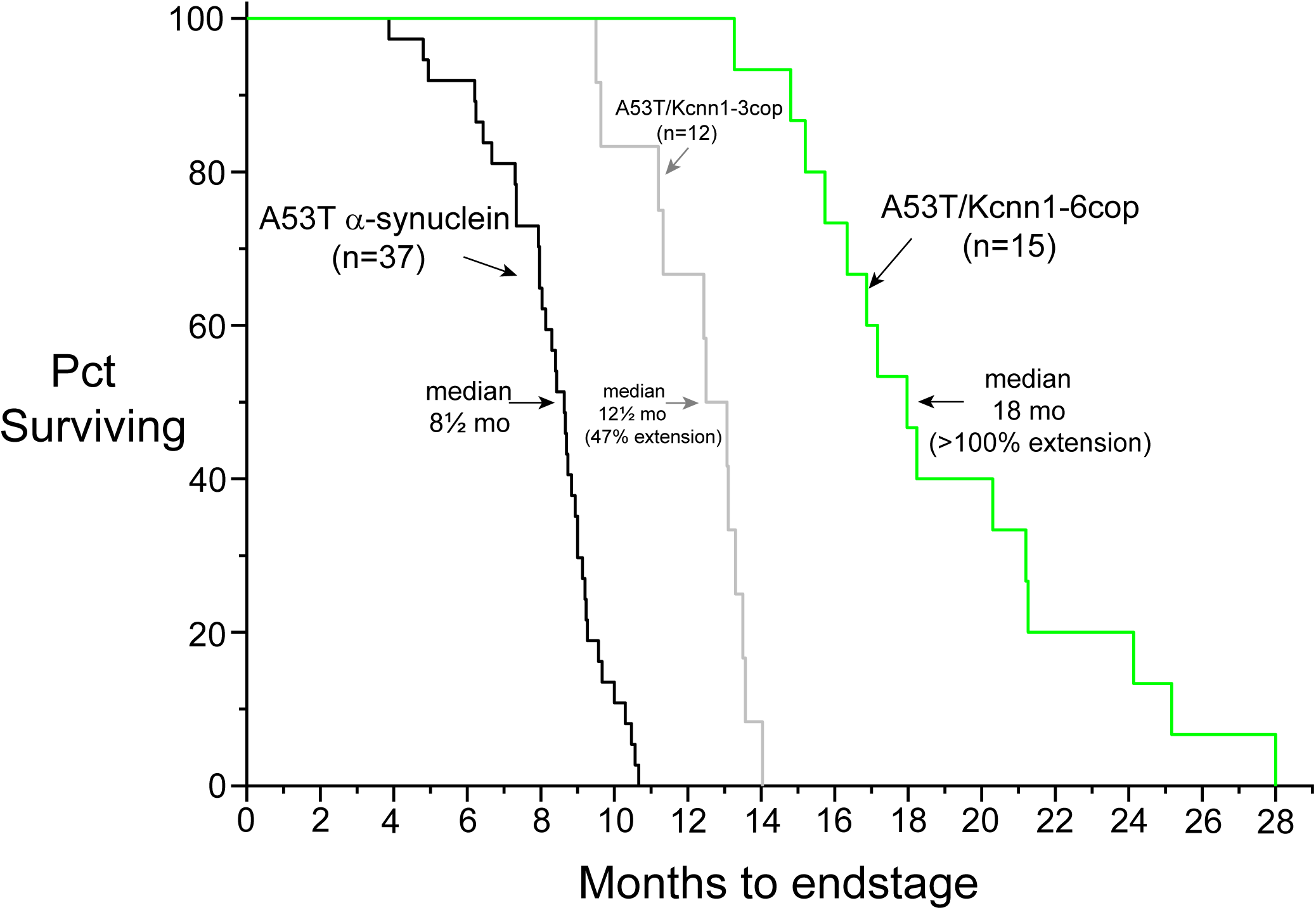
Kaplan-Meier survival plots for A53T α-synuclein transgenic mice, A53T/Kcnn1-3copy mice, and A53T/Kcnn1-6copy mice. Median survival time was extended by 47% for the A53T/ Kcnn1-3copy mice and >100% for the A53T/Kcnn1-6copy mice. The two Kcnn1 transgenes are independent. See text for the different motor phenotypes of the A53T and A53T/Kcnn1-6copy mice.

### Motor symptoms commence later in A53T/Kcnn1-6copy and differ from A53T alone

Corresponding to the longer survival of the A53T/Kcnn1-6copy mice was later onset of clinical disease, observed first at 12-16 mo of age. Differing from the A53T mice, which first exhibited forward flexion/splaying of the lower extremities at ∼7 months of age, the first abnormality of A53T/Kcnn1-6copy was clasping of a lower extremity when picked up by the tail. Initially, a short walk relieved this behavior. However, the elicited clasping progressed over a period of months to become bilateral. Even though walking was largely normal, at this point we initiated daily placement of wet food in a paper cup on the cage floor (performed also for A53T mice with splayed extremities). In some of the mice there was, by this time, a slowness to initiate movement, e.g. when prodded from behind (bradykinesia; ref.15). There was also in many cases an inability to build a nest, apparently a function of upper extremity weakness, indicated by poor upper extremity grasp of objects and poor wire hang performance (data not shown). At endstage, lower extremity clasping was accompanied by upper extremity clasping. At this stage, the mice preferred to lie on side, but could initially right, walk, and feed, but within a week or two they could no longer right and were sacrificed. Thus the nature of specific symptoms appeared to be different between A53T mice, where forward-flexed splayed lower extremities, a wobbling gait with sudden lurches and ultimately dystonic straight limb posture occurred, and A53T/Kcnn1-6copy mice, where symptoms were confined to clasping behavior (see Discussion). We conclude that Kcnn1 overexpression significantly altered both the time of onset and nature of the symptoms exhibited by the A53T mice.

### Alpha-synuclein disease-associated phospho-serine 129 α-synuclein is absent from A53T/Kcnn1-6copy mice at 12 months and 21 months as compared with A53T endstage mice

The extended survival of A53T/Kcnn1-6copy mice was associated with absence of accretion of phospho-serine 129 α-synuclein (hereafter, Psynuclein). This was shown first by sacrificing an asymptomatic A53T/Kcnn1-6copy mouse at 12 months of age, a time beyond the survival of any of the A53T mice but before symptom development for most of the A53T/Kcnn1-6copy mice. A sagittal brain section of the 12 month old A53T/Kcnn1-6copy mouse was compared with one from a 9 month old endstage A53T mouse, the two prepared in parallel with anti-Psynuclein immunostaining, and imaged with identical settings (Supp.Fig.1A,B). As shown in panel A, the end-stage A53T mouse exhibits strong Psynuclein staining in the same regions as previously reported^19^. In contrast, the 12 month old A53T/Kcnn1-6copy mouse shows no appreciable anti-Psynuclein signal, suggesting that the transgenic Kcnn1-6copy is conferring protection from α-synuclein toxicity in these regions.

As an example of putative Kcnn1-6copy protection, the superior colliculus, a midbrain region staining heavily with anti-Psynuclein in A53T endstage mice (circled in Supp.Fig.1A) is shown in Fig.2. This region, composed of a right and left superior colliculus, is involved with translating sensory inputs, particularly visual inputs (which enter superficial layers), to processing and motor outputs governing, e.g., head orientation (from intermediate and deep layers; see location of superior colliculi in Fig.2A sagittal and coronal schematics)^20,21^. Notably, in humans with diffuse Lewy body dementia, the superior colliculus was affected by α-synuclein, and there was loss of neurons from the intermediate gray layer (ref.22, Fig.3 therein). Here, in sagittal view of the superior colliculus of an endstage A53T mouse (9 mo.), there was copious Psynuclein staining (Fig.2B, top left panel), excepting for its superficial (visual input) region. By contrast, in a parallel reaction under the identical immunostaining and imaging conditions, the 12 month old A53T/Kcnn1-6copy mouse was devoid of Psynuclein immunostaining (Fig.2B, bottom left panel). The absence of Psynuclein corresponds with transgenic overexpression of Kcnn1 in the superior colliculus (bottom right panel; compare with endogenous level of Kcnn1 in absence of the transgene, top right panel). In the setting of overexpression (bottom right panel), a particularly prominent anti-Kcnn1 signal was observed in cells in what is likely to be the intermediate gray layer of the superior colliculus (arrowed). Thus, the absence of Psynuclein in the superior colliculus of the 12 month old A53T/Kcnn1-6copy mouse was associated with overexpression of Kcnn1 in the same region, supporting a protective role.

**Figure 2.**
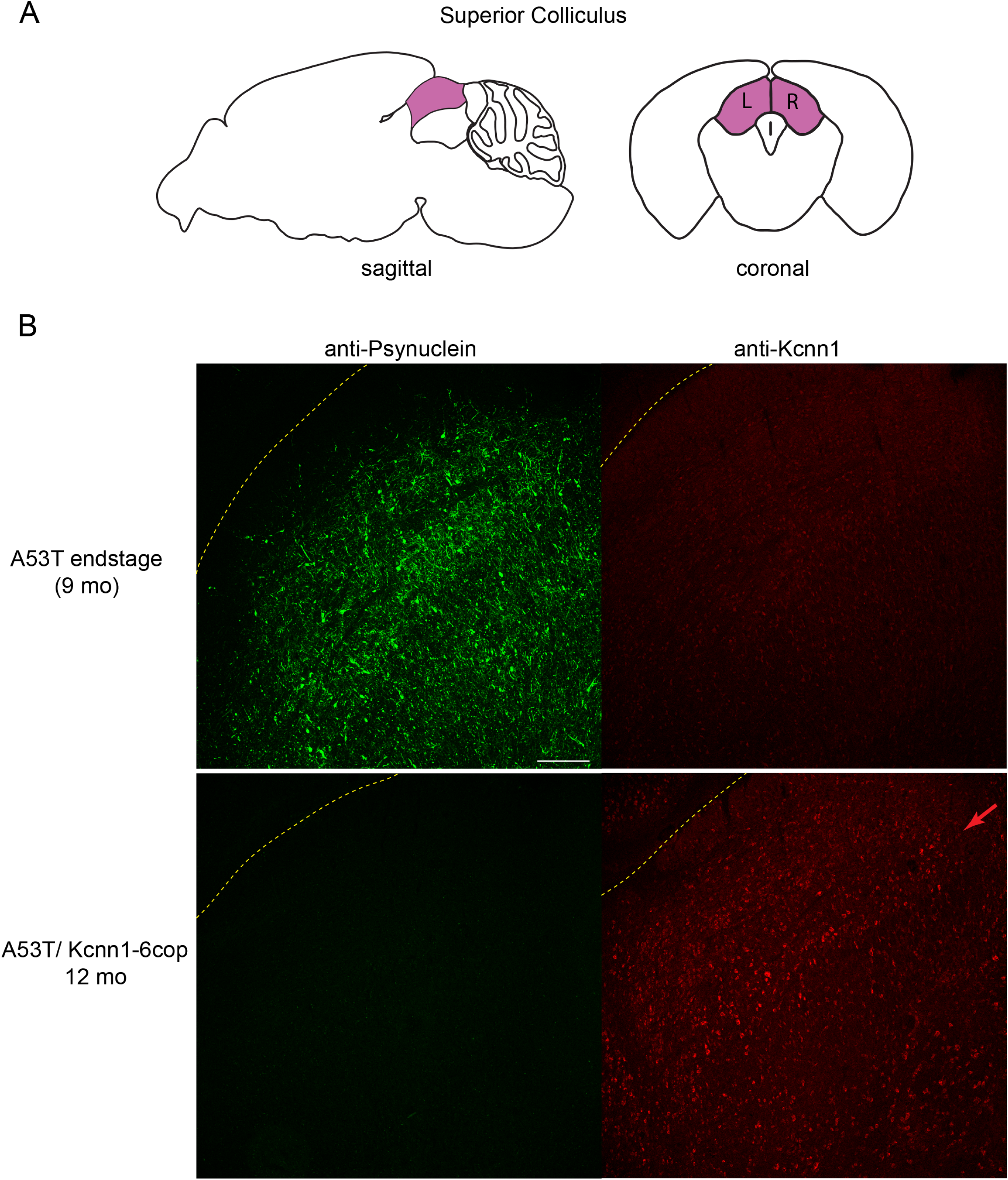
Absence of serine129 phospho-α-synuclein (Psynuclein) immunostaining in 12 mo A53T/Kcnn1-6copy mouse as compared with A53T endstage (9 mo) mouse. A. Location of superior colliculus (purple). Sagittal and coronal cross-sections of brain are shown, to illustrate midbrain dorsal position in the sagittal section and paired left and right structures in the coronal views. B. Sagittal views of superior colliculus, from 20 µm sagittal sections ∼1.2 mm from midline, showing presence of Kcnn1 overexpression in A53T/Kcnn1-6copy 12 month old mouse is associated with absence of α-synuclein disease-associated Psynuclein. Upper panels, A53T endstage 9 month old mouse sagitally sectioned and immunostained to reveal copious Psynuclein (left) and endogenous levels of Kcnn1 (right). Lower panels, absence of Psynuclein associated with overexpression of Kcnn1. Red arrow likely denotes intermediate gray layer. Scale bar, 200 μm.

**Figure 3.**
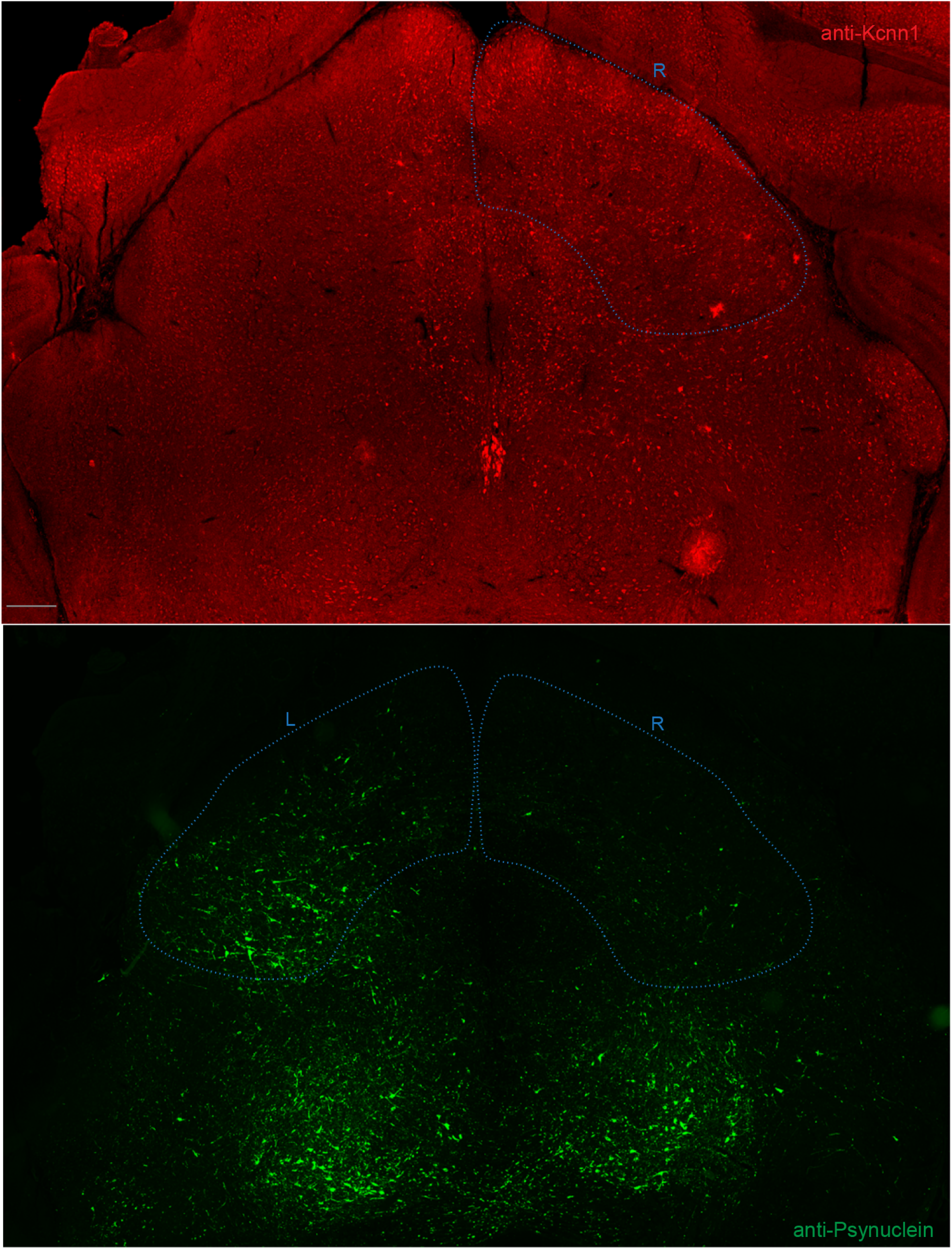
Stereotactic injection of AAV9-CMV-Kcnn1 virus into the right superior colliculus of an asymptomatic 6 month old A53T mouse leads to overexpression of Kcnn1 in right superior colliculus (upper panel, outlined in light blue), associated with absence of Psynuclein (lower panel outlined in blue) observed after 54 days. Shown in upper and lower panels are separately immunostained neighboring 20 µm cryosections of flash-frozen brain. Upper panel, coronal section of brain showing right superior colliculus with clusters of fluorescent anti-Kcnn1-positive cells in what appear to be the bands of superficial, intermediate, and deep gray layers. Fluorescent cells are also observed beyond the boundaries of the superior colliculus, including crossing the midline and ventrally (see text). Lower panel, almost complete absence of Psynuclein staining in injected right superior colliculus region vs non-injected left (outlined and labeled in blue). (For detail on Psynuclein accumulation outside superior colliculus, see text). Scale bar 250 μm.

More generally, examination of Kcnn1 expression in a sagittal section of the 12 month old A53T/Kcnn1-6 mouse (Supp.Fig.1D) showed many regions of overexpression in addition to superior colliculus, including e.g. zona incerta (ZI), thalamus (TH), hippocampal formation (HPF), deep cerebellar nuclei (DCN), vestibular nuclei (Ve), substantia nigra (SN), and cortex (compare with endogenous level of Kcnn1 expression from A53T alone endstage sagittal section (Supp.Fig.1C), prepared and examined in parallel. Among the Kcnn1-overexpressing regions (Supp.Fig.1D) were the same ones observed to accrete Psynuclein in absence of overexpression in A53T (circled in yellow). Thus, suppression of accretion of Psynuclein associated with overexpression of Kcnn1 in all of the regions where it accretes in the absence of Kcnn1 overexpression (A53T endstage panel A) indicates a broad ability of Kcnn1 overexpression to protect from production of the disease-associated Psynuclein.

The foregoing result with a 12 month old A53T/Kcnn1-6 copy mouse raised the question of whether there is also temporal breadth to the protection exerted by overexpression of Kcnn1, e.g. including later times when the symptom of lower extremity clasping when lifted by the tail is exhibited. A 21 month old mouse with lower extremity clasping was sacrificed and its brain was sagittally sectioned, stained with anti-Psynuclein, and imaged in parallel with a sagittal section from an A53T endstage mouse. As shown in Supp. Fig.1E, here also, no Psynuclein was detected as compared with the A53T mouse (Supp. Fig.1F). It thus appears that even at later times, in symptomatic A53T/Kcnn1-6 copy mice, disease-associated Psynuclein is not formed. Further studies of endstage mice, clasping all four extremities and unable to right, should address whether any Psynuclein is formed at the end-stage. Preliminarily, a detailed coronal analysis of one such mouse showed only rare flecks of Psynuclein in the regions floridly affected in A53T.

### AAV9-CMV-Kcnn1 virus injected into right superior colliculus of A53T mice at 6 months of age prevents Psynuclein formation in that location, associated with Kcnn1-overexpression

The transgenic mice just described overexpress Kcnn1 from early postnatal life, at the time when the Thy1.2 promoter becomes active. To assess whether effects on Psynuclein accumulation could be exerted by overexpressing mouse Kcnn1 in adult life, we stereotactically injected a self-complementary AAV9-CMV-mouse Kcnn1 virus (titer 6 x 10^13^vp/ml) into the superior colliculus in the right hemisphere of four presymptomatic A53T mice at 6 months of age (bregma −3.4 mm, lateral right 0.75 mm, 1.6 mm down from pial surface). We did not expect this site of injection to affect survival time of the mice and indeed the injected mice proceeded to develop the same progression of clinical disease to endstage as uninjected A53T mice. For immunofluorescence analysis at endstage, ∼150 consecutive 20 micron coronal brain sections were prepared from a flash frozen brain, starting rostral to the superior colliculi (at an arbitrary position ∼300-500 microns away; see sagittal view in Fig.2A) and proceeding in a rostral-caudal direction (essentially, from left to right in the sagittal view of Fig.2A). Every tenth coronal section was probed with anti-Psynuclein antibodies, and a neighboring section was probed with anti-Kcnn1 antibody.

Fig.3 shows the immunostaining results of two adjoining coronal sections (#59 and #60) from a region containing the superior colliculi of a mouse that had been injected on the right with Kcnn1 virus at 6 months of age and reached endstage 54 days later. Remarkably, there is nearly complete absence of Psynuclein (green) from the right superior colliculus (lower panel, outlined in blue), associated with overexpression of Kcnn1 (red) in the corresponding region (upper panel; outlined in blue). Note that there is broader expression of Kcnn1 beyond the right superior colliculus, extending across the midline to the left, and ventrally into the periaqueductal gray and mesencephalic reticular formation, as well as presence in the right medial geniculate (auditory relay nucleus; see lateral right zone) and the two Edinger Westphal nuclei (pupillary control nuclei; midline ventral to aqueduct). The former contiguous extended expression presumably results from local diffusion of injected virus, whereas the medial geniculate staining may result from viral tropism for this structure. The strong staining of EDW could be a result of viral tropism or, perhaps, strong endogenous expression.

In the lower panel of Fig.3 there is strong presence of Psynuclein in the left superior colliculus as compared with its nearly complete absence on the right. There is also bilateral Psynuclein in ventral structures including the mesencephalic reticular formation (eye movement control as well as other functions), and likely also in the red nucleus (motor coordination).(Note again that superficial layers of the superior colliculus do not accrete Psynuclein in A53T mice).

The same absence of Psynuclein from the injected right superior colliculus was observed throughout the coronally-sectioned body of the superior colliculus. Fig.4A shows a series of every tenth coronal section of sections from #49 to #99, stained for anti-Psynuclein (each separated by 200 microns, totalling ∼1000 microns of length). This revealed nearly complete absence of Psynuclein in the injected right superior colliculus as compared with its strong presence on the left.

**Figure 4AB.**
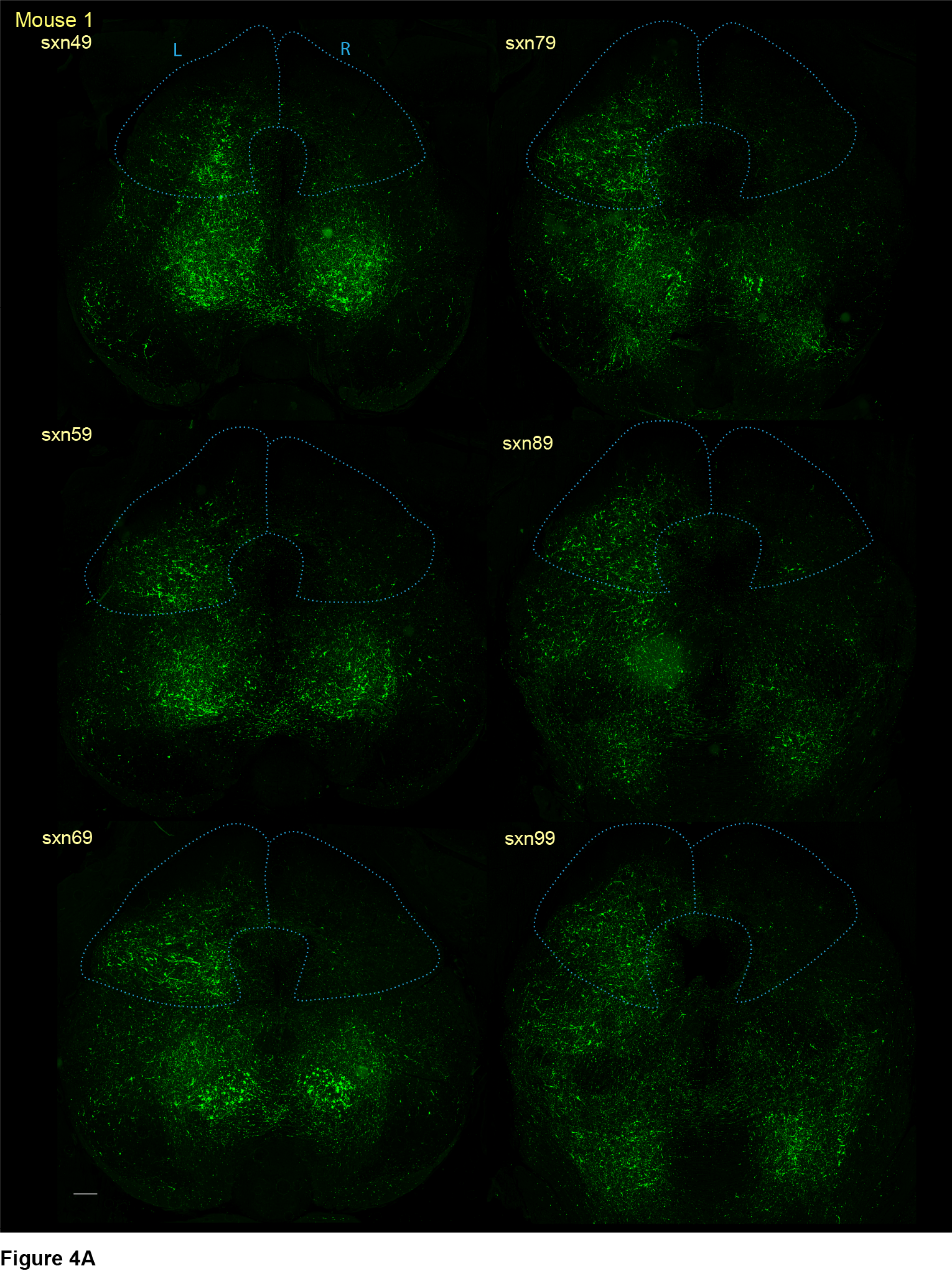
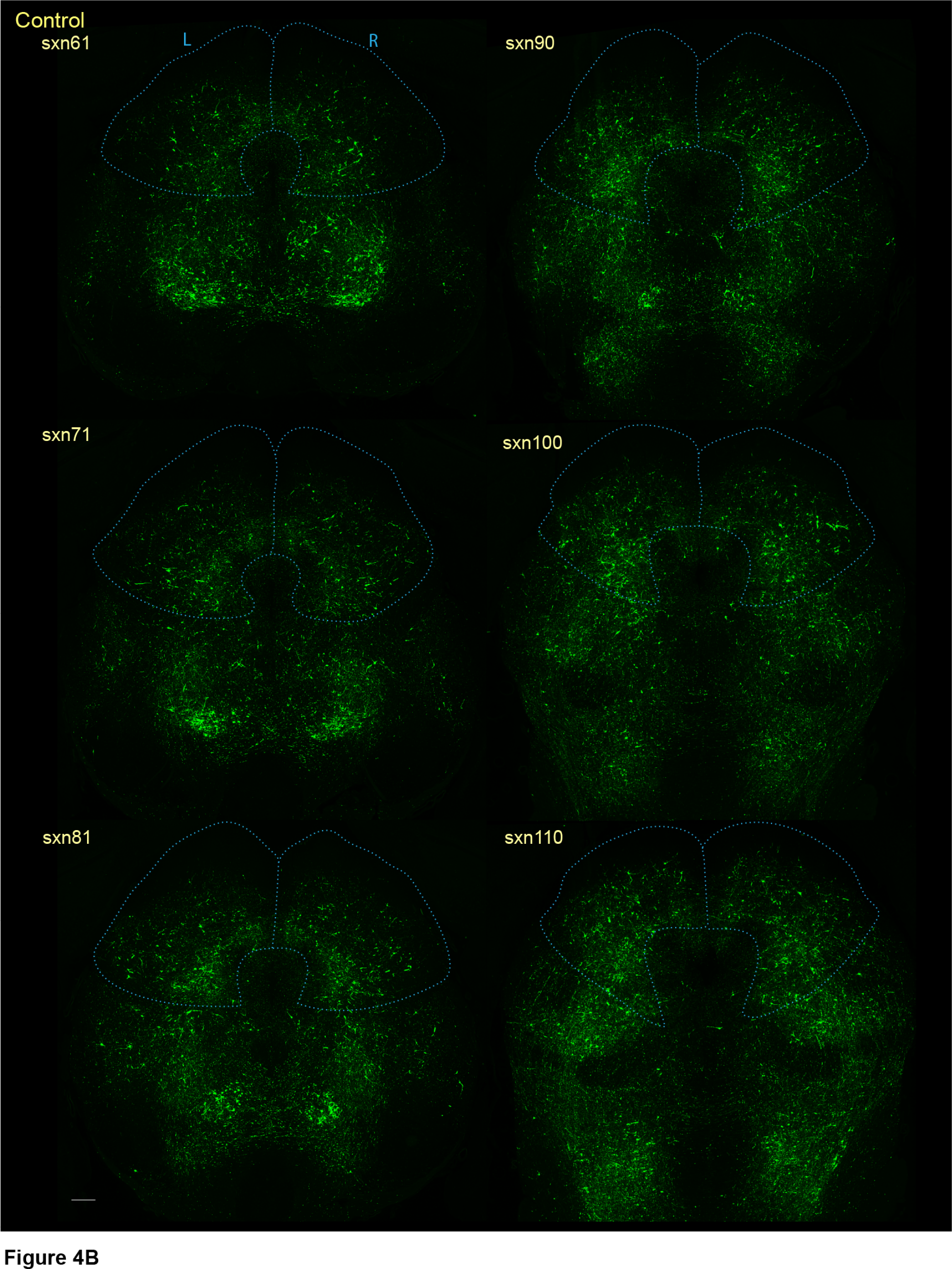
Absence of Psynuclein from right superior colliculus along ∼1 mm of rostro-caudal length of the AAV9-CMV-Kcnn1-injected mouse shown in Fig.3 (Mouse 1). The right and left superior colliculus are outlined in blue in all sections. Note that the shape of the outline is slightly altered from frame to frame to account for its position relative to surrounding landmarks (not shown) and compared with The Mouse Brain in Stereotaxic Coordinates, Franklin and Paxinos, 2007^32^. Coronal sections shown are spaced every 200 µm rostrocaudally. Consecutive 20 µm coronal sections of brain were taken, beginning at an arbitrary point several 100 µm rostral to superior colliculus and proceeding into the superior colliculus. Every 10^th^ section (sxn) was stained for Psynuclein. There is absence of Psynuclein from the injected right superior colliculus along the entire cut region. Section #89 appears to contain an artefact in its left aspect. Scale bar 250 μm. Figure 4B. A53T mouse injected with injection solution lacking virus (Control). Sections were collected and immunostained as in A. Superior colliculi circled in blue. There is symmetric presence of Psynuclein in the superior colliculi and outside of it. Scale bar 250 μm.

As controls, three mice were injected at 6 months with viral injection solution into the right superior colliculus and, upon reaching endstage and carrying out similar processing, all three exhibited copious bilateral Psynuclein in both the left and right superior colliculus (see Fig 4B, representative Control mouse, every tenth section, sections #61-#110).

The three other A53T mice, injected with Kcnn1 virus at 6 months of age, also exhibited reduction or absence of Psynuclein from right superior colliculus when similarly analyzed at endstage, but shorter rostro-caudal extent of effect was observed. For example, Psynuclein in Mouse 2 right superior colliculus, every tenth section from #43 to #93, is shown in Supp.Fig.2. Here, in section #43, Psynuclein was only modestly diminished, while more caudal sections were strongly diminished. By contrast, more rostral sections contained copious Psynuclein. The variability of suppression of Psynuclein is most likely a function of slightly different injection pathways from mouse to mouse, or differing diffusion paths of the injected virus from mouse to mouse. Consistent with Kcnn1 overexpression as nonetheless related to the effect, we observed that the regions with absence of Psynuclein correlated with sections with substantial Kcnn1 transduction (e.g. see Kcnn1 expression of section #64 of Mouse 2, the neighbor of Section #63, in Supp.Fig.3).

### Magnified views of left and right superior colliculi of AAV9 Kcnn1-injected A53T mouse at endstage

Magnified coronal views were obtained of the left and right superior colliculus of AAV9 CMV-Kcnn1 virus-injected Mouse 1 immunostained with anti-Psynuclein and anti-Kcnn1 (Fig.5). The magnified views are 400 micron x 400 micron squares taken with their centers measuring ∼1100 microns from either side of the midline at the level of the intermediate gray layer of the superior colliculus. In the uninjected left side (left panels), in the setting of an endogenous level of Kcnn1 (upper left panel), a dense amount of Psynuclein is observed inside both what appear to be neuronal cells and processes extending from them (lower left panel). Additional smaller amounts of Psynuclein may be present in glia or the neuropil. In contrast, on the virus-injected right side, intermediate gray layer somata, likely neuronal, are more heavily immunostained by anti-Kcnn1 (upper right panel). Whether glia are also immunostained could not be resolved. Resembling earlier Thy1.2-Kcnn1 transgenic studies (Fig.2 and ref.19), viral-transduced Kcnn1 is excluded from the nucleus and is present at a level of stained protein that is 5-10 times the level of endogenous protein. Associated with Kcnn1 overexpression on the virus-injected right side, there was nearly complete absence of Psynuclein (right lower panel; a single cell soma appears to be immunostained as well as 4-5 “flecks”). Thus Kcnn1 overexpression in neurons of the superior colliculus, including the intermediate gray layer, appears to prevent the local formation of Psynuclein. The absence of Psynuclein was not a consequence of loss of neurons on the right side, as shown by Nissl staining, where no apparent loss of cells or architecture or staining was observed (not shown); by counting Kcnn1-positive cells from the lateral left (after increasing brightness) and lateral right intermediate gray layers (in 600 μm X 600 μm squares), where no difference was observed (data not shown); and by costaining with anti-Kcnn1 and anti-NeuN (Supp.Fig.5), which shows on both left and right that Kcnn1 cells are NeuN-positive, and that they are roughly equal in number on both left and right sides. The results overall suggest that one or more steps of the process of α-synuclein toxicity that is associated with accumulation of Psynuclein is prevented by overexpression of Kcnn1. This correlates overall with protection from clinical disease and enables extended survival observed with the Thy1.2-Kcnn1 transgene expression in A53T (Fig.1).

**Figure 5.**
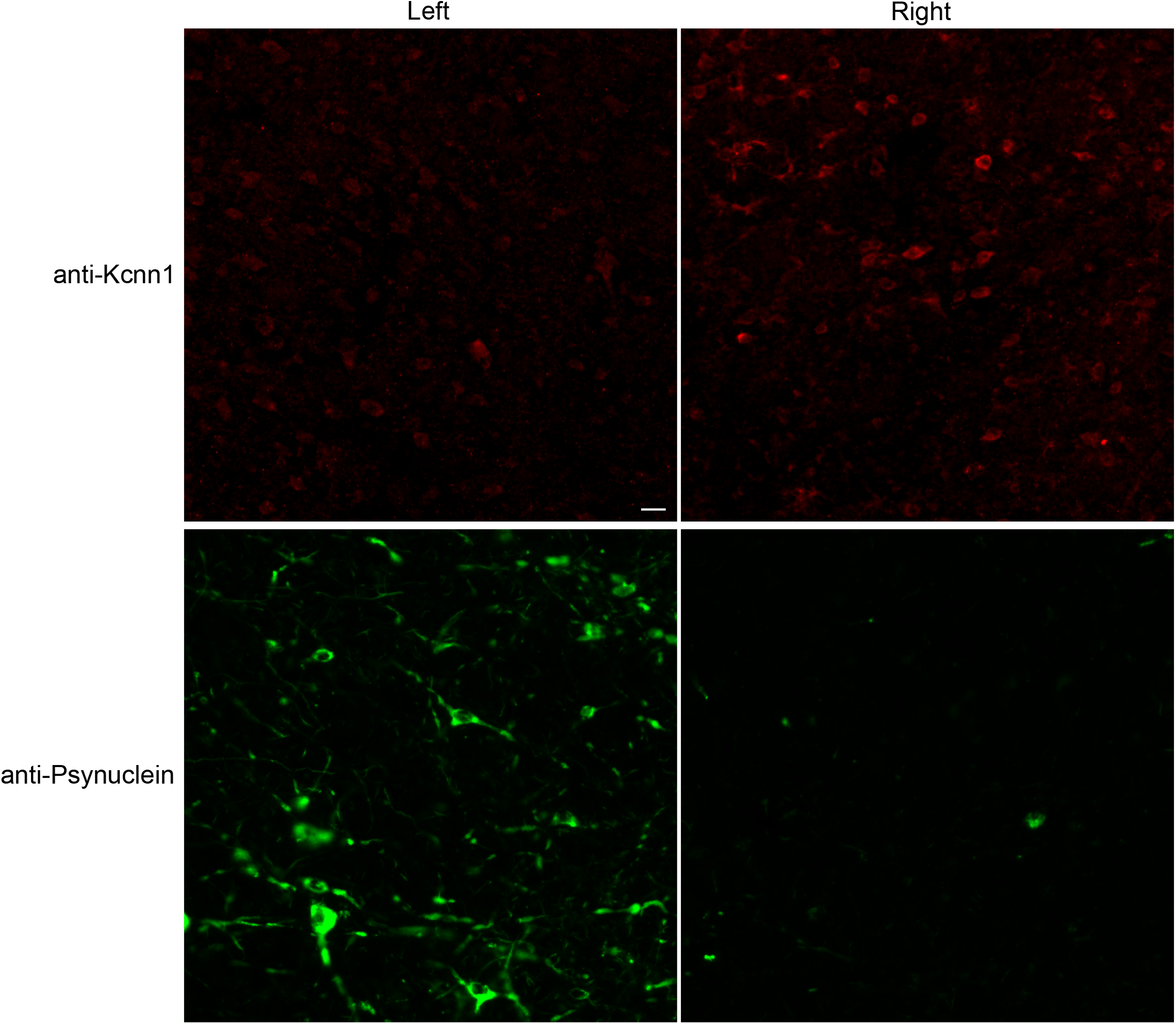
Magnified views of corresponding regions of left and right superior colliculus from an AAV9 CMV-Kcnn1 virus-injected mouse at level of intermediate gray layers, immunostained with anti-Kcnn1 and anti-P-synuclein, showing near complete absence of Psynuclein in the presence of overexpressed Kcnn1. Displayed are 400 µm X 400 µm squares, taken from coronal section #60 (Kcnn1) and section #59 (Psynuclein) of Mouse 1, with their centers ∼1100 microns from either side of the midline. Anti-Kcnn1 (upper panels) shows brighter Kcnn1-immunoreactive cells on the virus-injected right side vs much less bright cells (endogenously expressing Kcnn1) on the uninjected left side. The number of Kcnn1-immunostained cells was roughly equal on the two sides (data not shown). Anti-Psynuclein (lower panels) shows copious intracellular Psynuclein on the left, in the setting of endogenous Kcnn1, and also that Psynuclein extends into processes and that a portion may be in glia or neuropil. On the right, in the setting of overexpressed Kcnn1, only four or five flecks of Psynuclein are visible, the rightmost of which appears to be intracellular. Scale bar 20 µm.

## Discussion

In two settings involving a transgenic A53T mutant α-synuclein mouse strain with fully penetrant and lethal motor disease, overexpression of Kcnn1 appears to prevent the formation of a disease-associated form of α-synuclein, phospho-serine 129 α-synuclein (Psynuclein). The first setting involves the constitutive expression of Kcnn1 from a Thy1.2-Kcnn1 transgene that extended median survival from 8.5 months to 18 months (Fig.1). Examination of a 12 month old A53T/Kcnn1-6copy mouse (asymptomatic) revealed overexpression of Kcnn1 in neurons of the superior colliculus, but no accretion of Psynuclein (Fig.2). This compared to abundant accumulation of Psynuclein in the superior colliculus of endstage A53T mice (Fig.2 and Supp Fig.1A). This protective behavior was more general, extending to other regions of A53T endstage mice that had accumulated Psynuclein (Supp.Fig.1A), where overexpression of Kcnn1 (Supp.Fig.1D) likewise suppressed its formation (Supp.Fig.1B). Prevention of Psynuclein formation also extended temporally to a symptomatic timepoint (21 mo) of an A53T/Kcnn1-6 mouse (Supp.Fig.1E). The second setting involved injection of a self-complementary AAV9 CMV-Kcnn1 virus into the right superior colliculus of a presymptomatic adult A53T mouse (6 mo), which showed overexpression of Kcnn1 in the right superior colliculus (but not the left superior colliculus) two months later at endstage, associated with near complete absence of Psynuclein in the right superior colliculus (Figs.3 and 4). Questions remain as to how the observed effects are exerted.

### Possible mechanisms of beneficial effects of Kcnn1 overexpression

One possibility is that Kcnn1 might be protecting in a cell autonomous manner the Thy1.2-Kcnn1-overexpressing neurons from toxicity from the A53T α-synuclein. An earlier study indicated that Kcnn1 overexpression is associated with an ER stress response involving the UPR and ISR^19^. These may offer neuroprotection in this setting, resembling the effect of such Kcnn1 overexpression to protect spinal cord motor neurons in a G85R SOD1YFP-linked ALS mouse model from aggregation of the transgenic mutant SOD1-YFP protein, correlated there with increased median survival^19^. We find overexpressed Kcnn1 to be localized to the ER membrane (unpublished observations), but in this location it may have additional effects to that reported earlier of producing ER stress^19^, such as channel function. Whether such effects are mediated at the neuronal soma or synapse are unknown, but notably, Psynuclein could be found in both soma and neurites in an A53T endstage mouse (Fig.5 left lower panel as an example). Effects of overexpression of Kcnn1 to prevent toxicity of misfolded/aggregating α-synuclein could involve stimulating autophagy, e.g. via activating ERphagy through expansion of the ER as was previously observed^19^.

An alternative possibility is that Kcnn1 overexpression begets a cellular response of microglia, astrocytes, or oligodendrocytes that acts to block toxicity from A53T α-synuclein. Concerning potential microglial removal of neurons containing Psynuclein, we did not observe loss of neurons in the AAV9-Kcnn1-injected right superior colliculus (Supp.Fig.4).

### Modification of clinical phenotype

Notably, in the setting of constitutive transgenic overexpression of (Thy1.2)-Kcnn1, the A53T α-synuclein-driven motor disease is modified by Kcnn1 from dystonic posturing of extremities (of A53T alone) to a relatively slowly progressive extremity clasping that does not commence until 12-16 months of age. Thus, among the specific sites of Kcnn1 overexpression, one or more is exerting a strong effect on the clinical manifestation of α-synuclein toxicity. The most immediately observable histologic correlate in the A53T/Kcnn1 transgenic mice is the prevention of accumulation of disease-associated Psynuclein.

Whether the clasping behavior is simply a milder form of the dystonia exhibited by the A53T mice or is a separate motor pathology is unclear. LaLonde and Strazielle^23^ have pointed out that any of forebrain, cerebellum, basal ganglia, midbrain, or spinal cord can be implicated in clasping behavior, and that 10% of B6SJL mice (used here) display clasping, imputed to the SJL strain contribution. Yet dystonic behavior in mice also can be associated with any or some combination of the indicated structures (e.g. refs. 24-28). Thus the nature of the Kcnn1-modified behavior remains to be resolved. It is not purely a Kcnn1 transgenic-related symptom, however, because the Kcnn1-6copy mice do not exhibit clasping at any age.

Finally, it should be noted that various dystonic symptoms have been reported in patients with several early-onset forms of Parkinson’s Disease, including those with mutations in *PARKIN, PINK, DJ-1,* and in idiopathic patients on L-DOPA therapy during “off” periods^29^, but the relationship to the A53T mouse clinical behavior and modification reported here remains unclear. Concerning the L-DOPA-related behavior, we have no evidence for effects on the SNpc in A53T mice: the SNpc is minimally stained by anti-Psynuclein and the number of TH-positive cells does not differ between SNpc of A53T and B6SJL mice (data not shown). Yet the effects of Kcnn1 overexpression to suppress accumulation of Psynuclein in many brain nuclei would suggest that it could have beneficial effects in the nigrostriatum if α-synuclein-driven pathology were present.

### Can Kcnn1 virus reverse α-synuclein toxicity in addition to preventing it?

A further question concerns whether Kcnn1 virus is able to reverse pre-existing Psynuclein-associated disease processes, beyond what appears from the present data to be a preventive action. It seems difficult to resolve reversibility without a serial examination of tissue, which is not feasible with the need here to acquire tissue for immunostaining. While virus transduction clearly shows effects at the level of presymptomatic adult animals, it would be unclear, if symptomatic mice were virus-injected, how to distinguish, at the level of a single mouse, the reversal of pathology vs simply forestalling of additional pathology before endstage. Other approaches, e.g. observing an improved behavioral output, seem needed.

## Acknowledgments

This work was supported by a Yale Sterling Endowment and by AWM Bio Inc. We thank Yale Animal Resources Center for excellent technical assistance. We thank J.Cardin for technical advice on stereotactic injection. We thank M.Guerra and Confocal Microscopy at CCMI and S.Wilson at Imaging Core Facility in Neuroscience for superb technical assistance in image acquisition. A.L.H., W.A.F., and M.N. are co-founders of AWM Bio, Inc. which is the exclusive licensee of certain pending patent applications covering this work.

## METHODS

### Mice

All animal experiments were carried out under protocols approved by the Yale University Institutional Animal Care and Use Committee in accordance with National Institutes of Health guidelines for the ethical treatment of animals. Wild-type B6SJLF1/J mice (JAX strain #100012), called B6SJL, were purchased from Jackson Laboratories. The hemizygous Thy1-Kcnn1 3-copy and 6-copy transgenic strains were described previously^19^, as were mice hemizygous for a transgene of Thy1.2-driven human A53T α-synuclein cDNA^6^ that were maintained on a mixed B6SJL background^19^. The genotypes of all animals were confirmed by quantitative real-time PCR using primers suggested by Primer-BLAST from NCBI (www.ncbi.nlm.nih.gov/tools/primer-blast/).

### Viruses

A plasmid (pscAAV-GFP, Addgene #32396, a gift from John T. Gray, St. Jude Children’s Research Hospital, Memphis, TN)^30^ designed to produce self-complementary AAV-CMVGFP virus was modified by direct replacement of the GFP sequence with one encoding mouse Kcnn1 cDNA. The mouse Kcnn1 cDNA sequence (NM_032397), was PCR-amplified from OriGene Technologies plasmid MR208601, omitting the 3’-terminal Myc-DDK tag sequence. The final construct was sequenced after preparation. This plasmid was packaged at the University of North Carolina Chapel Hill Vector Core to produce recombinant self-complementary (sc) AAV virus with the AAV9 serotype at a titer of 6.6 x 10^13^ vp/ml.

### Stereotactic injection of virus

Stereotactic injection of the AAV9 scCMV-Kcnn1 virus was carried out following the surgical procedure detailed in ref #31. A Digital Mouse Stereotaxic Instrument (Stoelting), including stage, injector, and motorized injection controller, was used. Coordinates for injection were: AP = 0.4 mm; ML = +0.75 mm; DV = −1.6 mm. A single burr hole was drilled in the right side of the skull at those AP and ML coordinates using a 0.5 mm bit (Meisinger US#1/4) attached to a hand-held electric nail drill. The dura was pierced with a 27G needle. A 5 µl Hamilton syringe (848511) was lowered to the DV coordinate and then used to deliver 1.0 µl of the virus-containing solution via a 34G injection needle (Hamilton 207434) at a rate of 0.1 µl/min, followed by slow retraction.

### Antibodies

Antibodies used: rabbit anti-mouse Kcnn1 (Proteintech 17929-1-AP, 1:400 dilution), rabbit anti-phospho (S129) α-synuclein (Abcam ab51253,1:800), anti-NeuN (Rbfox3)(Millipore MAB377, 1:100). Secondary antibodies, Alexa Fluor 488 donkey anti-rabbit, Alexa Fluor 568 donkey anti-rabbit, Alexa Fluor 488 Goat anti-rabbit and Alexa Fluor 568 goat anti-mouse IgG1, were purchased from Invitrogen and used at 1:500 dilution.

### Immunostaining

Because successful staining with the Kcnn1 antibody required using freshly dissected (unfixed) tissue, mice undergoing immunochemical analysis were transcardially perfused with PBS alone. After dissection, brain segments were immediately embedded in OCT (Tissue-Tek) and frozen on LN-chilled methyl butane, then stored at −80°C. Twenty µm coronal or sagittal sections of brain were prepared on a cryostat (Leica CM3050S), placed on Surgipath X-tra slides, and stored at −80°C. For immunostaining, slides were post-fixed for 25 minutes in 4% paraformaldehyde in PBS, then washed 3 times in PBS. Sections were permeabilized with 0.5% Triton X-100 for 20 min, then blocked with buffer containing 0.3 M glycine, 1% BSA, 5% goat or donkey serum, 0.1% Triton X-100 in PBS. Primary antibodies were diluted into the blocking buffer lacking glycine and incubated for 3 hr room temperature, then overnight at 4 C. After PBS washes the secondary antibody was applied for 3 hr at room temperature. After additional washing steps slides were mounted with Vectashield containing DAPI (Vector Labs).

### Image Acquisition

Images were acquired on a Leica SP5 confocal microscope or an Olympus VS200 Slide Scanner using a 10x objective. Excitation and emission parameters appropriate for AlexaFluor 488, AlexaFluor 568, and DAPI were used, depending on the microscope and the fluor examined. Leica LAS-X software was used to display and export the images from the Leica microscopes, and OlyVIA was used to process the images from the Olympus scope, in both cases generating TIF files for producing illustrations. ImageJ software was used to pseudocolor images, and Adobe Photoshop was used to adjust the brightness and contrast of images. Where there are multiple images in individual figures, all have been adjusted to identical brightness/contrast values. Images were assembled in Adobe Illustrator.

