## Supplemental Figure 1 for "Neuronal overexpression of mouse potassium channel subunit Kcnn1 in A53T α-synuclein mice more than doubles median survival time, associated with suppression of phospho-S129 α-synuclein formation" (from another version of this preprint: version 1, 2026-03-11)

A

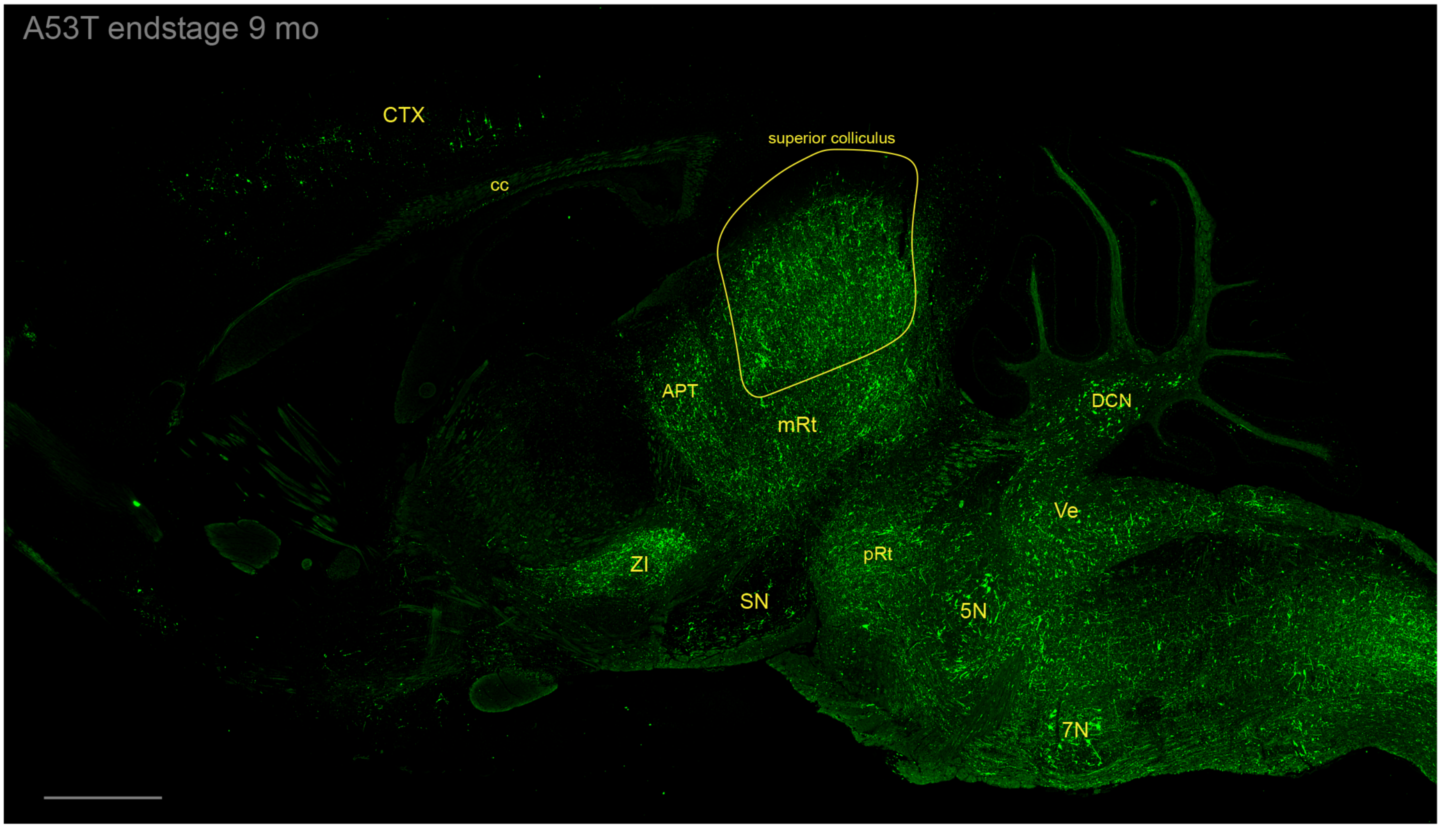


B

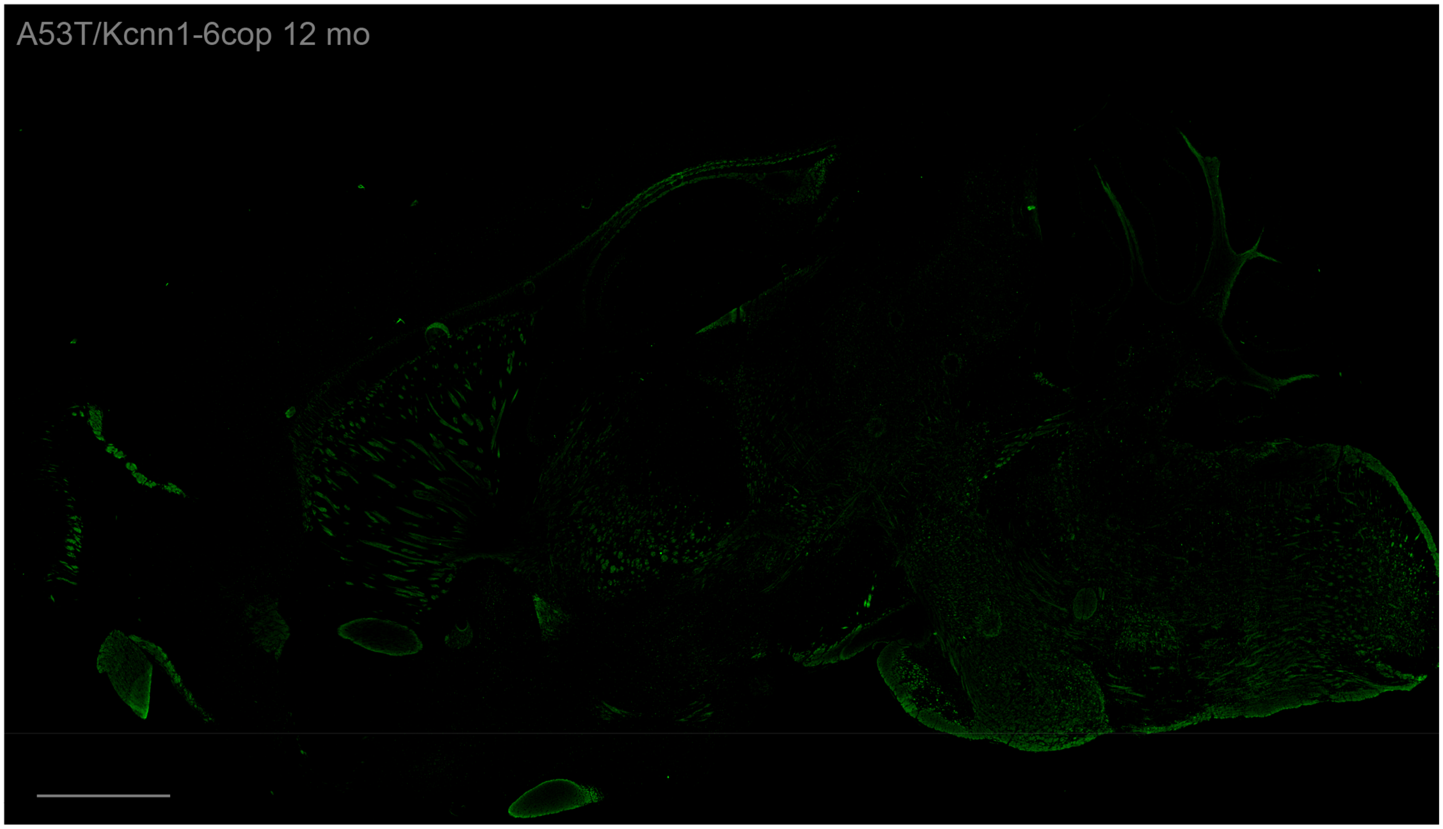


**Supp. Fig.1 A,B**

C

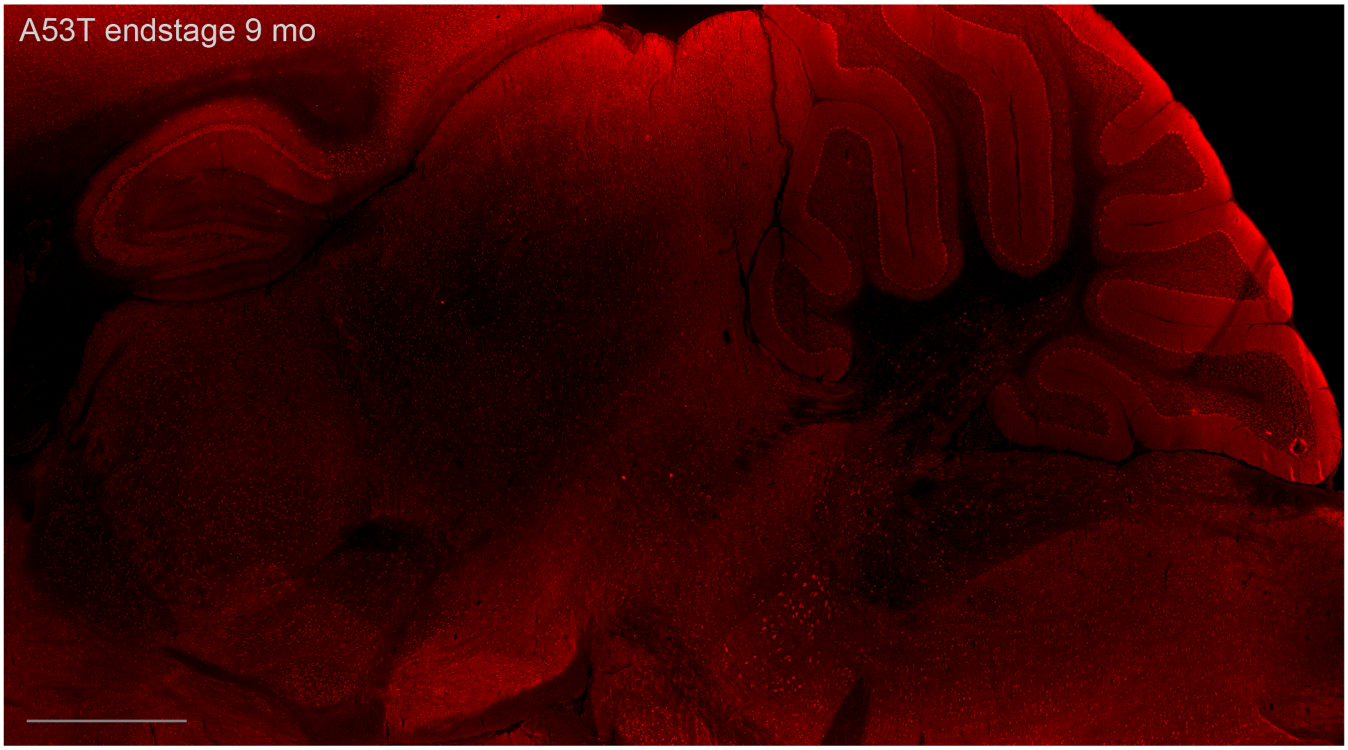


D

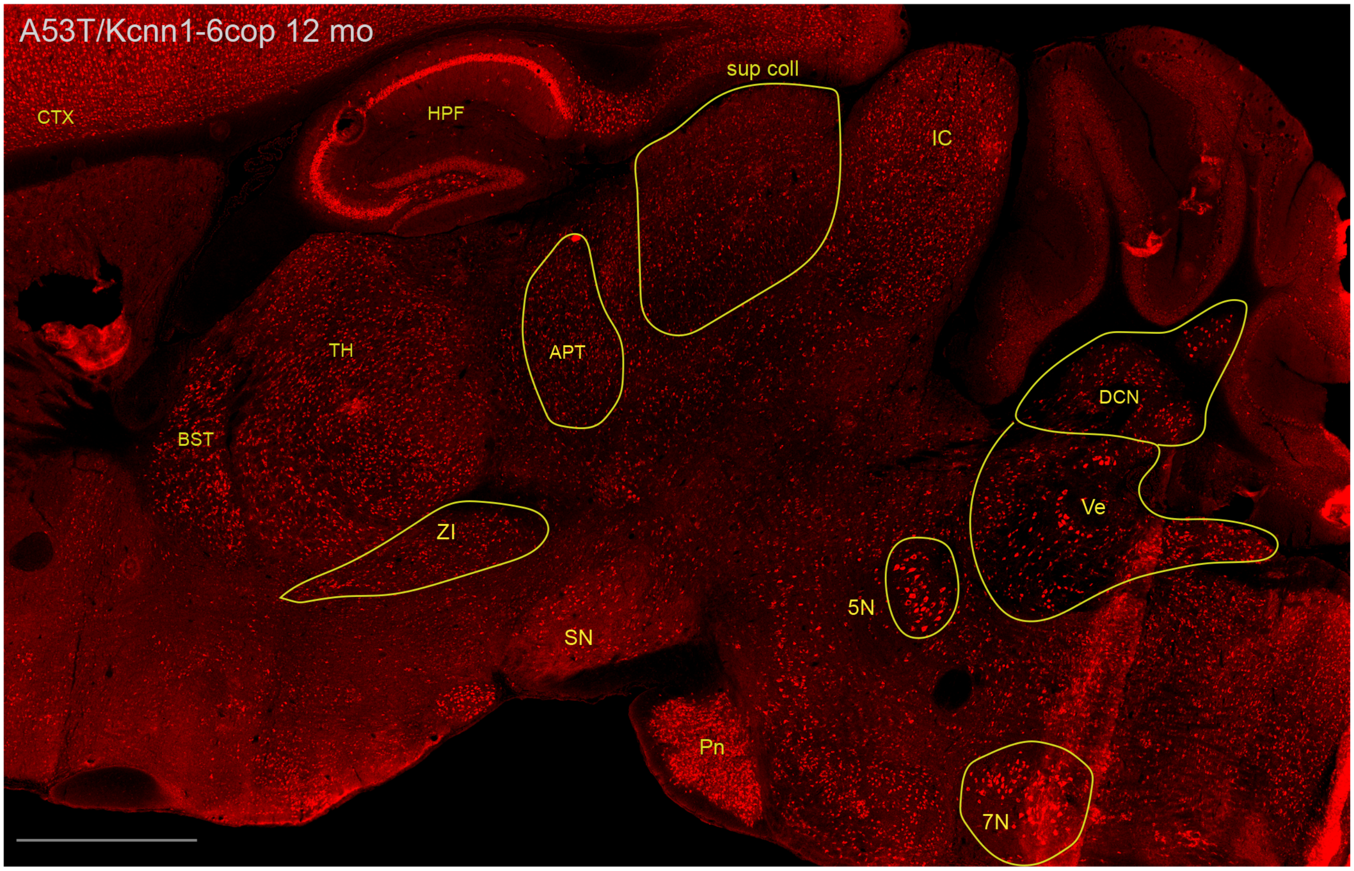


**Supp. Fig.1 C,D**

E

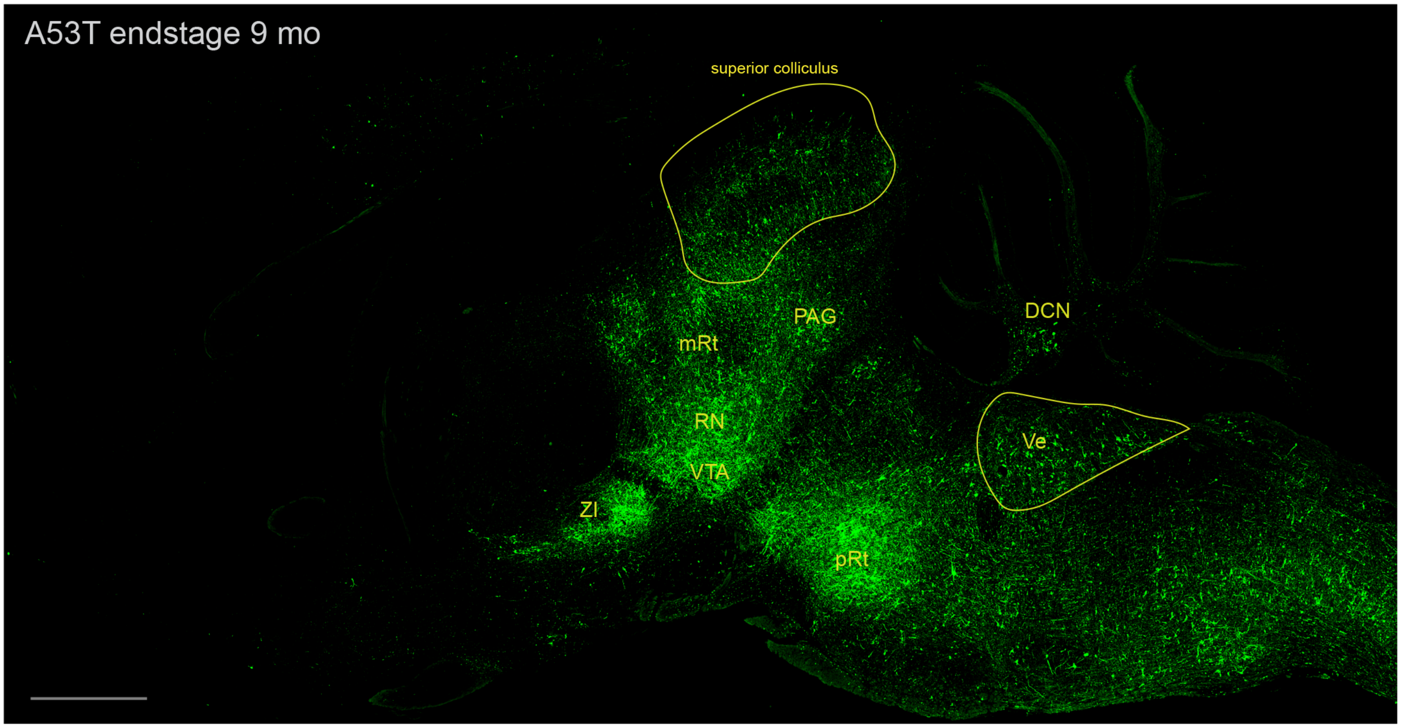


F

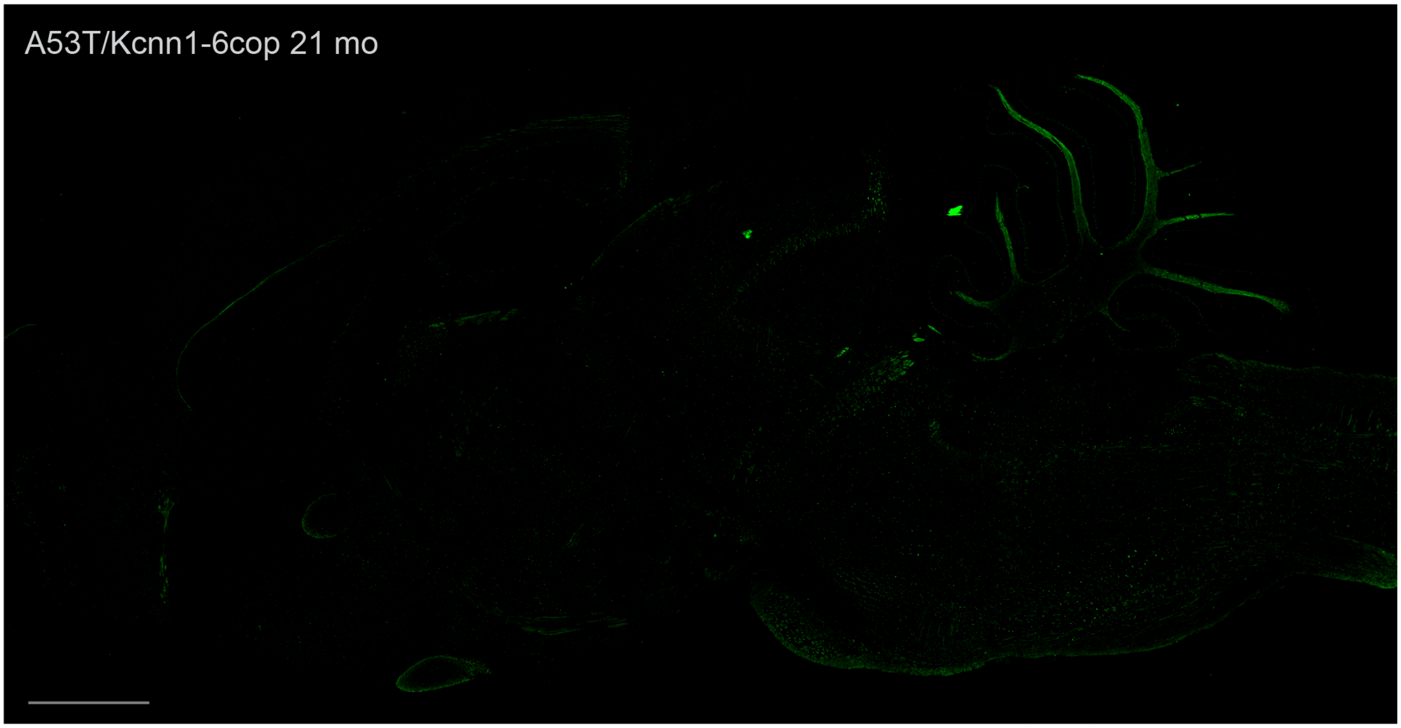


**Supp Fig. 1.A.** Strong anti-serine129-Phosphosynuclein immunostaining of superior colliculus and other regions as shown in a whole brain sagittal section (1.3 mm from midline) of an endstage A53T-α-synuclein transgenic mouse at 9 months of age. Freshly frozen brain was sagittally sectioned, then immunostained with anti-Psynuclein antibody. Superior colliculus in midbrain region is outlined in yellow. Other regions that are also strongly immunostained are indicated, including ZI (zona incerta), 5N (trigeminal nucleus), DCN (deep cerebellar nuclei), and Ve (vestibular nuclei). Less strongly labeled are mRt (mesencephalic reticular formation) and APT (anterior-pretectal nucleus), as well as CTX (cortex). Minimally labeled is the SN (substantia nigra). cc, corpus callosum. Scale bar 1 mm.
**B.** No significant anti-serine129-Phosphosynuclein immunostaining of 12 month old A53T/Kcnn1-6copy transgenic mouse. Brain was prepared, sectioned, and a section (1.0 mm from the midline) immunostained as described in **A**.

**C.** Anti-Kcnn1 immunostaining of sagittal brain section of endstage A53T α-synuclein transgenic mouse shown in panel **A**, taken ~1.3 mm lateral to the midline, showing endogenous expression levels of Kcnn1. Endogenous expression apparently cannot suppress the formation of Psynuclein observed in A53T (panel **A**), including the trigeminal nucleus which weakly stains with endogenous Kcnn1. Note that the whole brain sagittal image has been cropped to minimize edge regions that stain heavily (see e.g. cerebellar surface), for reasons unknown.
**D.** Anti-Kcnn1 immunostaining of sagittal section of asymptomatic 12 moth old A53T/Kcnn1-6copy transgenic mouse, which is devoid of Psynuclein as shown in panel **B** (taken ~1.0 mm lateral to the midline, from the same immune staining reaction as **C**, and with the same imaging settings and with similar cropping of the figure as in panel **C**. Overexpression of Kcnn1 is noticeable in a broad range of structures including all of those, outlined and labeled, that exhibited strong Psynuclein in endstage A53T (see panel **A**), including 5N which had a low level of endogenous Kcnn1 in endstage A53T (panel **C**). Kcnn1 overexpression in the 12 month old A53T/Kcnn1-6copy mouse thus correlates with the lack of Psynuclein formation that is otherwise observed in endstage A53T mice, suggesting the protective action of Kcnn1 overexpression is general to a variety of neuronal nuclei. Additional regions, unaffected with Psynuclein in A53T endstage, also express abundant Kcnn1, including e.g. substantia nigra, thalamus, and inferior colliculus. Note that the portion of cortex in upper left of panel **D** that exhibited Psynuclein in Supp.Fig.1**A** is not demarcated. Note also that neighboring brain regions not shown in this image do not exhibit overexpression).

**E.** Absence of significant accumulation of Psynuclein in a 21 month old A53T/Kcnn1-6copy transgenic mouse symptomatic with bilateral lower extremity clasping. Immunostaining was carried out on a sagittal section 0.6 mm lateral to the midline in parallel with a sagittal section of an endstage A53T, shown in panel **F** for comparison.
