## Supplemental Figure 2 for "Neuronal overexpression of mouse potassium channel subunit Kcnn1 in A53T α-synuclein mice more than doubles median survival time, associated with suppression of phospho-S129 α-synuclein formation" (from another version of this preprint: version 1, 2026-03-11)

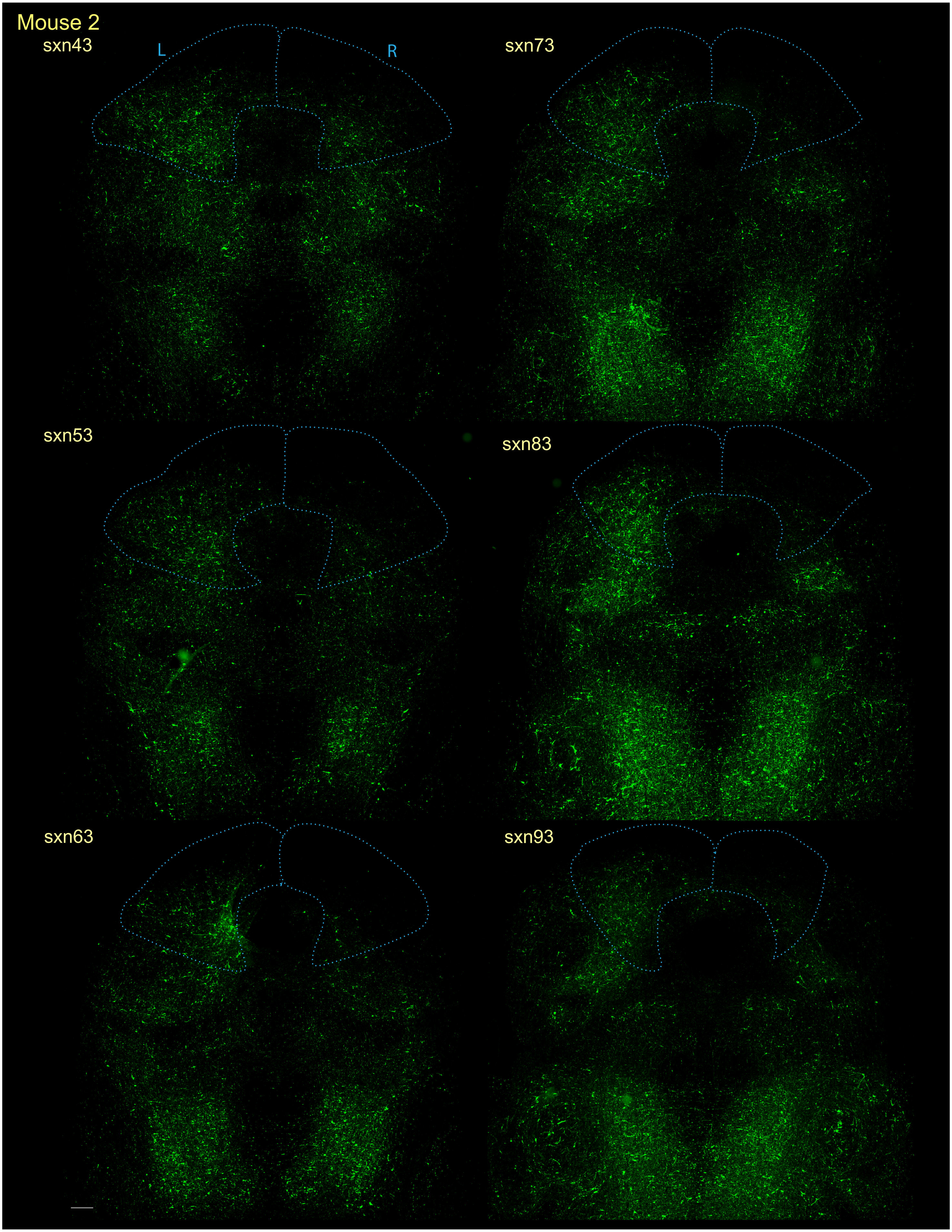


**Supp. Fig. 2.** Variability of Psynuclein absence from right superior colliculus following AAV9-CMV-Kcnn1 virus injection into superior colliculus at 6 months. This mouse (Mouse 2) was injected into the right superior colliculus while asymptomatic at 6 months and then analyzed at endstage (9 mo). Here, there is only partial absence of Psynuclein from the right superior colliculus in sxn #43 (and no absence rostrally, not shown). More complete absence is observed from sxn #53 to #93, comprising a rostral-caudal distance of ~800 εm. Anti-Psynuclein staining of other portions of the sections are as observed in other images and described in the text related to Fig.3. Scale bar 250 εm.
