## Supplemental Figure 3 for "Neuronal overexpression of mouse potassium channel subunit Kcnn1 in A53T α-synuclein mice more than doubles median survival time, associated with suppression of phospho-S129 α-synuclein formation" (from another version of this preprint: version 1, 2026-03-11)

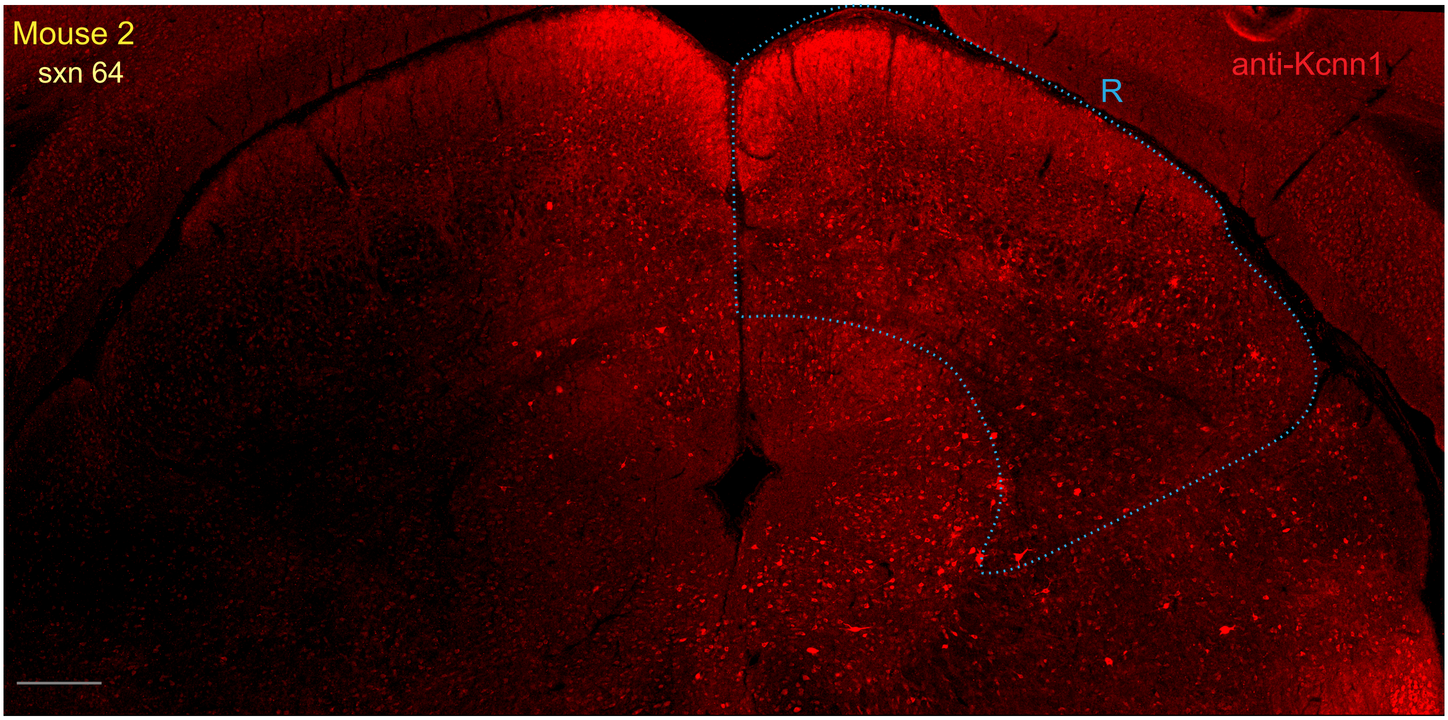


**Supp. Fig. 3.** Despite variability of absence of Psynuclein following AAV9-CMV-Kcnn1 injection, sections with right-sided absence of Psynuclein have corresponding overexpression of Kcnn1. Section #64 of Mouse 2, neighboring one with complete absence of Psynuclein in right superior colliculus (#63 in Supp. Fig. 2), is Kcnn1-positive in the right superior colliculus (outlined in blue). Brightly staining clusters of Kcnn1-positive cells are observed, as before, in layers that correspond to the gray layers. The nature of the very positive tissue staining at the most superficial layer remains unknown. Scale bar 250 μm.
