## Supplemental Figure 4 for "Neuronal overexpression of mouse potassium channel subunit Kcnn1 in A53T α-synuclein mice more than doubles median survival time, associated with suppression of phospho-S129 α-synuclein formation" (from another version of this preprint: version 1, 2026-03-11)

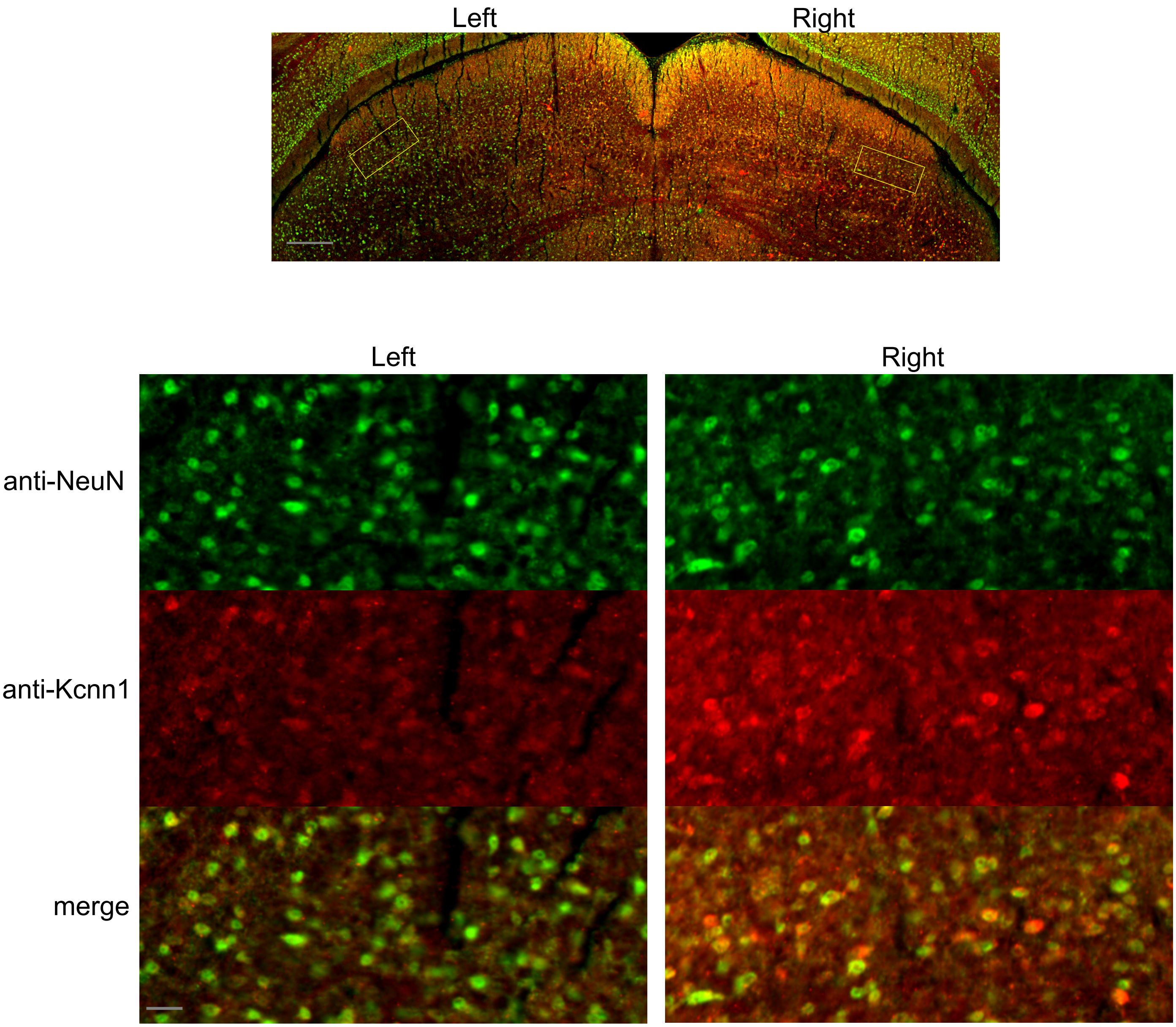


**Supp. Fig. 4.** No evidence that neuronal loss explains absence of Psynuclein from Kcnn1 virus-injected right superior colliculus.
Top panel, superior colliculus left and right regions, double-immunostained with anti-NeuN (green), marking neurons, and Kcnn1 (red), marking endogenous levels of Kcnn1 (left) and of overexpressed Kcnn1 in addition to endogenous (right). The orange color on the right results from merging of the green and red immunostaining colors. Scale bar 250 μm.
Lateral regions at the level of middle layers of the left and right superior colliculi in the top panel are boxed in yellow (280 μm X 120 μm) and displayed at high magnification below, showing immunostaining with anti-NeuN (top), anti-Kcnn1 (middle), and merging of the two (bottom). For anti-NeuN, the density and numbers of immunostaining neurons at left and right are roughly equal. For anti-Kcnn1, immunostaining on the left is relatively weak, comprising endogenous Kcnn1, while many of the Kcnn1-positive cells at the right are considerably brighter, likely comprising AAV-Kcnn1-transduced cells that appear to be mostly neurons, considering that they match up with cells in the NeuN panel at top right. The merge in the panel at lower left is overwhelmingly green, but the weakly anti-Kcnn1-positive cells in the middle panel match up well with neurons in the top left panel, suggesting that most endogenous Kcnn1-expressing cells are neurons. The merge in the panel at lower right directly supports such a matchup in the setting of Kcnn1 overexpression by showing many cells with an orange tint resulting from combination of green and red, while other cells display a green nucleus, where NeuN localizes, and red cytoplasm, where Kcnn1 localizes. Scale bar 20 μm. The green and red channels were merged in ImageJ to produce the top panel and the "merge" of the bottom panel.
